# Inherited mitochondrial dysfunction triggered by OPA1 mutation impacts the sensory innervation fibre identity, functionality and regenerative potential in the cornea

**DOI:** 10.1101/2024.04.18.590110

**Authors:** Léna Meneux, Nadège Feret, Sarah Pernot, Mélissa Girard, Solange Sarkis, Alicia Caballero Megido, Mélanie Quiles, Laura Fichter, Jerome Viaralet, Christophe Hirtz, Cécile Delettre, Frederic Michon

## Abstract

Mitochondrial dysfunctions are detrimental to organ metabolism. The cornea, transparent outmost layer of the eye, is prone to environmental aggressions, such as UV light, and therefore dependent on adequate mitochondrial function. While several reports have linked corneal defects to mitochondrial dysfunction, the impact of OPA1 mutation, known to induce such dysfunction, has never been studied in this context. We used the mouse line carrying OPA1^delTTAG^ mutation to investigate its impact on corneal biology. To our surprise, neither the tear film composition nor the corneal epithelial transcriptomic signature were altered upon OPA1 mutation. However, when analyzing the corneal innervation, we discovered an undersensitivity of the cornea upon the mutation, but an increased innervation volume at 3 months. Furthermore, the fibre identity changed with a decrease of the SP+ axons. Finally, we demonstrated that the innervation regeneration was less efficient and less functional in OPA1^+/-^ corneas. Altogether, our study describes the resilience of the corneal epithelial biology, reflecting the mitohormesis induced by the OPA1 mutation, and the adaptation of the corneal innervation to maintain its functionality despite its morphogenesis defects. These findings will participate to a better understanding of the mitochondrial dysfunction on peripheral innervation.

## Introduction

Mitochondrial dysfunctions gather all conditions or diseases due to an abnormal number of mitochondria, a dysfunction in the respiratory chain, or the incapacity to provide the substrate for mitochondrial metabolism^1^. These defects can be either inherited or non-inherited. The latter can occur due to spontaneous mtDNA mutations or somatic mutations that accumulate with age, rendering the mitochondrial metabolism less efficient with time^2^. On the other hand, inherited mitochondrial dysfunctions result from mutations in mitochondrial DNA (mtDNA) or nuclear gene defects affecting mitochondrial function^3^. Among the known pathologies originating from an inherited mitochondrial dysfunction, the dominant optic atrophy (DOA) is an inherited blinding disease caused by the degenescence of the retinal cell ganglions forming the optic nerve^4^, some patients declare as well extra-ocular syndromes such as ataxia, deafness and peripheral neuropathies^5,6^. Mutations in the *OPA1* gene are responsible for the majority of the DOA diagnostics. *OPA1* gene encodes a mitochondrial dynamin like GTPase involved in the fusion process and is essential to maintain the organite integrity. It was recently shown that different OPA1 mutations lead to distinct mitochondrial structural impariments^7^. A mouse model, bearing the OPA1^delTTAG^ mutation, recapitulates the degenerative syndromes encountered in DOA patients^6^. Furthermore, this mouse model exhibits multi-systemic phenotype, such as ataxia, cardiomyopathy and peripheral neuropathy. Finally, it was shown that this mutation leads to mitochondrial supercomplex instability, from which cell death originates^6^.

While the impact of mitochondrial dysfunctions on the posterior segment of the eye (retina and optic nerve) has been heavily documented, few reports have linked these dysfunctions to pathologies affecting the anterior segment (cornea, iris and lens). Inherited mitochondrial dysfunction is a key factor in the progression of Fuchs endothelial corneal dystrophy (FECD) and keratoconus^8–10^. Case reports have linked mitochondrial diseases to corneal cloudiness^11,12^, perforation^13^ and edema^14^. Furthermore, mitochondrial defect was associated to inflammatory context and ROS production in ocular surface pathologies^8,15^. While mitochondrial dysfunction is associated with lens cloudiness^11,12^, no report has linked DOA to ocular surface defects. Despite these studies, the direct effect of mitochondrial disfunction on corneal biology remains elusive.

Among the key structures of the anterior segment, the cornea is the transparent tissue covering the eye. Its main roles are to convey the light toward the retina, and to act as a protective barrier against infection or injury for the eye. The transparency of the cornea is a crucial component of its function and is the result of specific architecture and cell arrangements. The corneal structure is composed of a monolayered endothelium on the posterior side, on which the stroma lies. This part is a thick layer of extracellular matrix with a low cell density^16–19^. Finally the anterior layer is a life-long renewed pluristratified epithelium^20^.

To coordinate epithelial homeostasis and healing, and to adequately respond to any change in the corneal environment, this tissue is highly innervated^21,22^. The corneal innervation is mainly composed of Aδ and C sensory fibres whose cell bodies are located in the ophthalmic branch of the trigeminal ganglion^23^. Those fibres penetrate the corneal stroma from the periphery, as bundles of fibres branching and spreading within the stroma^24^. When entering in the epithelium, the fibres branche again and radiate horizontally to form the subbasal plexus. Finally, the sensory fibres shift orientation to reach the top of the epithelium, where they end up as free nerve endings^21,22^. Stimuli transduction is not the only role of the corneal innervation, as these fibres have a crucial role on epithelial homeostasis through the secretion of trophic factors essential for epithelial cell survival^25^.

Due to its localization, the cornea is highly exposed to ultraviolet radiation and high oxygen tension, making it particularly vulnerable to mitochondrial defects^8^. Moreover, the DOA patients exhibit peripheral neuropathy, which can have a dramatic effect on corneal epithelial homeostasis. Therefore, we investigated the impact of OPA1^delTTAG^ mutation on mouse corneal biology. In the current study, we found an unexpectedly healthy corneal epithelium upon OPA1 mutation. Given the role of OPA1 in the maintenance of mitochondria, and cornea being subjected to a heavy oxidative stress, we expected a progressive deleterious phenotype of OPA1 mutated cornea. However, we found a physiological tear proteome and normal corneal transcriptomic signature. Our analyses showed that upon OPA1 mutation, the dense corneal innervation was altered. We identified a connection between OPA1 and corneal sensory fibre identity. Our data indicate a decorrelation between the identity of the fibres and their functionality. Mechanistically, we found a link between OPA1 and corneal innervation maturation.

## Results

### OPA1 mutation has no impact on the tear film

As the OPA1^delTTAG^ mutation was never studied in the corneal context, we first validated the impact of the mutation on *OPA1* corneal expression in the OPA1^+/-^ animals by qPCR (Fig.S1). We measured a close to 50% decrease of *OPA1* expression in the cornea of OPA1+/- animals, confirming the validity of this mutation in the tissue. To get a broader sense of mitochondrial dysfunction impact on corneal biology, we analyzed the corneal microenvironment. Among the crucial elements constituting the corneal microenvironment, we first investigated the tear film proteome composition. Therefore, we sampled the tear film from OPA1^+/+^ and OPA1^+/-^ mice. Interstingly, the OPA1^+/-^ mice displayed a similar tear production when compared to the littermate control animals at 3 and 6 months of age (Fig.1A). After assessing the quantity of tears, we used sensitive untargeted mass spectrometry to analyse the tear proteome quality. Among the identified proteins, we detected factors known to be present in tears, such as lactotransferrin^26^, serum albumin^27^, prolactin inducing protein^28^, peroxiredoxin 6 and gelsolin^29^, confirming the efficiency of the proteomic analysis. Similarly to the tear production, the proteomic signature was almost not altered in the context of OPA1 mutation, regardless of the animal age (Fig.1B). Only 3 proteins out of 266 showed a significant p-value modulation at 3 months, and 22 out of 269 at 6 months. However, these factors, i.e. keratin 19, annexin A4, ras-related protein rap1b, and acyl coA binding protein are intracellular proteins, probably originating from corneal epithelial cells shed in the tear film. Besides looking at the p-values, we used Benjamini–Hochberg p-value correction to analyse the q-values. While we obtained a more precise analysis of the tear film, it only confirmed the absence of modification of its composition, neither at 3 nor 6 months. Taken together, these results demonstrated the lack of impact of mitochondrial defect on tear film quality and quantity, reflecting the stability of the tear film, and the lacrimal gland integrity.

**Figure 1.**
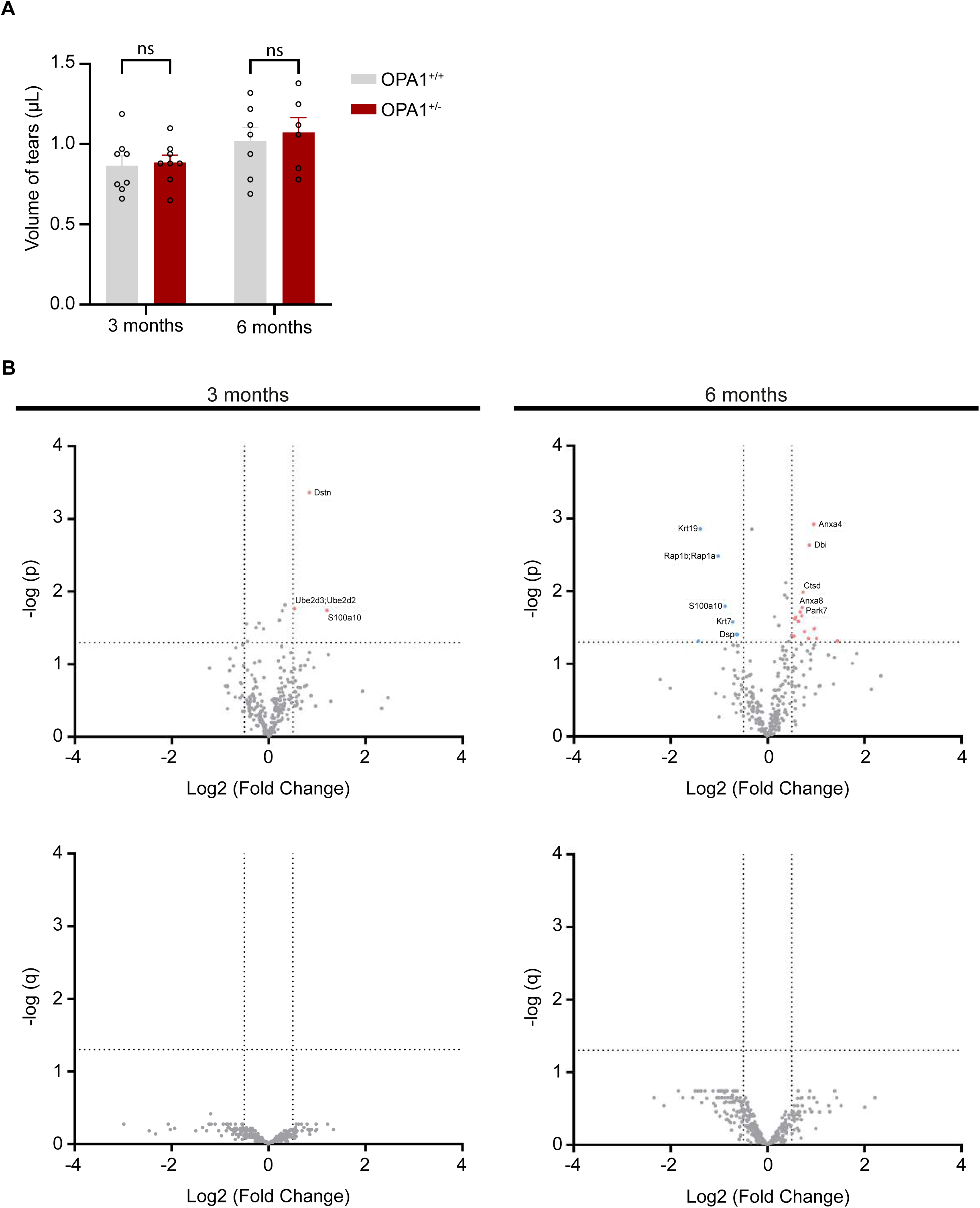
Tear film composition analysis reveals no modification upon OPA1 mutation. **(A)** Tears were collected to evaluate the tear production. Tear volume measurements are similar between OPA1^+/-^ and OPA1^+/+^ animals at 3 and 6 months (n=6-8 mice per group). Statistical analysis using two-way ANOVA with Sidak’s multiple comparisons test. Significant p-value are lower than or equal to 0.05. **(B)** Volcano plot representations of murine tear proteomic analysis by mass spectrometry (n=3 per group). The horizontal dotted lines represent the limit of statistical significance. The vertical dotted lines separate the identified proteins presenting a fold lower than 0.5 (blue dots), or higher than 2 (red dots). The identified proteins not satisfying these parameters are represented with grey dots. Top volcano plots are represented with p-values.Three proteins are presenting a fold over 2 at 3 months and 21 proteins are withing these parameters at 6 months. Statistical analysis using a t-test with permutated-based FDR of 0.05. Bottom volcano plots are represented with q-values. No protein are satisfying the parameters. Statistical analysis using a Benjamini-Hochberg p-value correction.

### The corneal epithelium is not altered upon OPA1 mutation

As corneal defects has been reported in the mitochondrial dysfunction context^30^, we next analyzed the effect of the OPA1 mutation on the corneal biology. First, we studied the corneal structure through the measurement of corneal thickness in OPA1^+/-^ animals, using optical coherence tomography (OCT) imaging. We assessed the full corneal, the epithelial and stromal thickness in 3 and 6 month-old animals (Fig.2). Our results did not detect any thickness alterations in OPA1 mutant mouse cornea. We confirmed this observation with Hematoxylin-Eosin-Safran histological staining (Fig. S2). Furthermore, the OCT and histological staining did not present any defected tissue organization.

**Figure 2.**
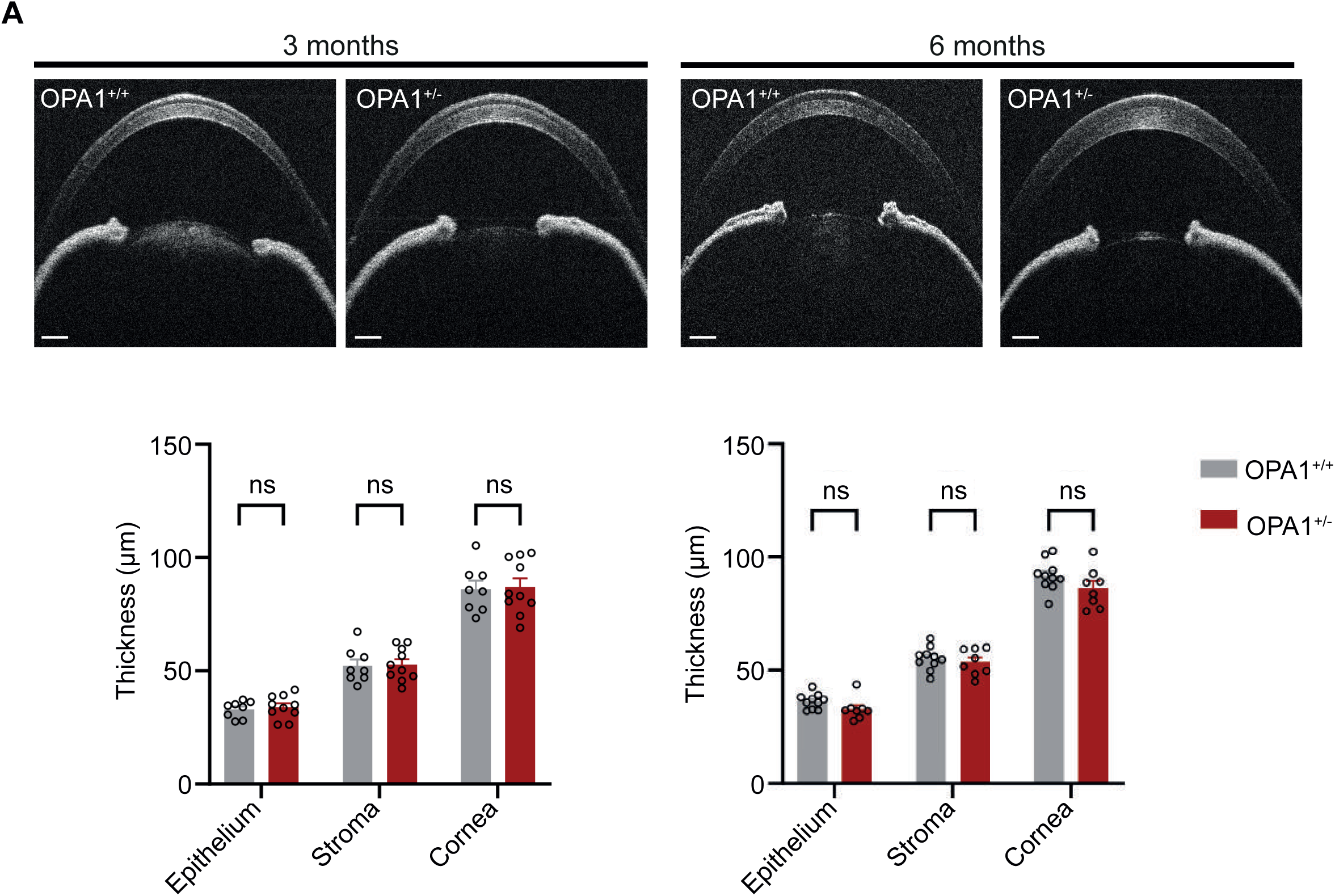
Corneal gross morphology of the OPA1^+/-^ mice is normal. Thicknesss measurements from SD-OCT images of the cornea and its layers in OPA1^+/+^ and OPA1^+/-^ animals at 3 and 6 months (n=8-10) highlight the similarity of the corneal structure between mutant and wildtype animals. Data are represented as mean ±SEM. Statistical analysis using two-way ANOVA with Sidak’s multiple comparisons test. Significant p-values are lower than or equal to 0.05.

Because of the lack of gross morphology modifications resulting from OPA1 mutation, we investigated possible alteration of the corneal transcriptomic signature, which could reflect more subtle changes in the cell metabolism or terminal differentiation (Fig.3). Using bulk RNA-sequencing on corneas from 3 and 6 month-old animals (Fig.3A), we found only 2 genes significantly misregulated in 3 month-old OPA1^+/-^ corneas (*Nxph4* was upregulated and *Alox15* was downregulated). Furthermore, in 6 month-old OPA1^+/-^ corneas, we identified 9 genes that were misregulated (6 upregulated and 3 downregulated). Despite the low number of differentially expressed genes, we analysed the expression of 8 of them via qPCR. We selected the 2 transcripts (*Nxph4* and *Alox15*) from 3 months, and 3 transcripts the most downregulated (*Adgrf2*, *Arntl*, and *Clmp*) and 3 the most upregulated (*Adamts1*, *Ciart*, and *Cpeb1*) at 6months. From these 8 candidates, only Cepb1 displayed a significant overexpression in the OPA1^+/-^ cornea (Fig.3B), only *Cpeb1* expression. As this factor has been involved in epithelial-to-mesenchyme transition ^31^, and no other transcripts were affected, it seemed that this level of upregulation was not enough to have a strong effect on corneal epithelial biology. We concluded that the OPA1 mutation had close to no effect on the corneal physiology.

**Figure 3.**
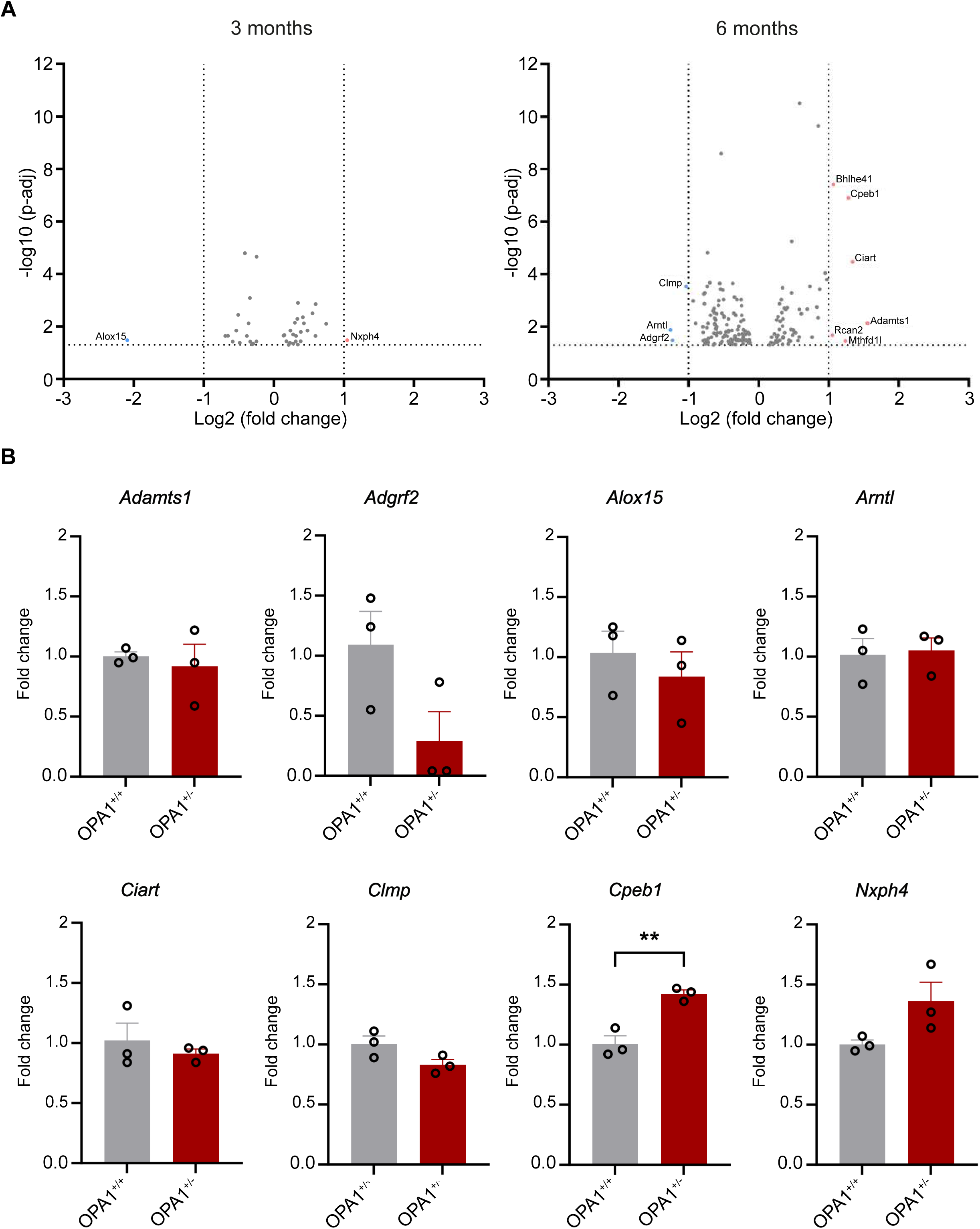
Transcriptomic corneal analysis analysis confirms the absence of major corneal modification. **(A)** The RNA-sequencing shows that the corneal transcriptomic signature is globally unaltered upon OPA1 mutation. The horizontal dotted lines represent the limit of statistical significance. The vertical dotted lines separate the identified transcripts presenting a fold lower than 0.5 (blue dots), or higher than 2 (red dots). At 3 months, 2 transcripts satisfy these parameters, and 9 at 6 months. The identified transcripts not satisfying these parameters are represented with grey dots. Significant p adjusted-values are lower than or equal to 0.05. **(B)** RT-qPCR analysis of the three proteins with the lowest and highest significant fold change. Only Cpeb1 confirm the results obtained in (A). Data are represented as mean ±SEM. Statistical anlaysis using unpaired t-test and, **p<0.01.

### Mitochondrial dysfunction from OPA1 mutation modifies the corneal innervation

After investigating the tear film and the corneal biology, we focused on the last element of the corneal microenvironment, its innervation. The dense corneal innervation secretes crucial trophic factors for the epithelium^22,32^, and is essential to sense any damages of the ocular surface^33,34^. Therefore, we studied the distribution of the nerve fibres and their functionality upon mitochondrial dysfunction. To evaluate the innervation density, we used a software to calculate the volume of stained fibres obtained with a pan neuronal antibody (anti βIII-tubulin) in the central cornea (Fig.4A). While we did not notice any sexual differences regarding the corneal gross mophology, , in the OPA1^+/+^ female mice, the volume of innervation increased between 3 and 6 months of age, contrasting with OPA1^+/-^ animals exhibiting a steady innervation volume from 3 to 6 months (Fig.4B). Furthermore, 3 month-old OPA1^+/-^ females displayed a significantly greater volume of innervation, reflecting a maximal volume of innervation reached earlier in life (Fig.4A-B). Notably, the OPA1^+/-^ males did not exhibit any volume innvervation defects at 3 months (Fig.S3), therefore, we decided to focus the rest of our study on OPA1^+/-^ females. Then, we tested the innervation functionality, using Von Frey filaments, similarly to using a Cochet-Bonnet aesthesiometer on patients^35,36^. Despite a higher volume of innervation at 3 months, the OPA1^+/-^ females showed a lower sensitivity than in OPA1^+/+^ females. This discrepancy was not anymore found in 6 month-old animals (Fig.4C).

**Figure 4.**
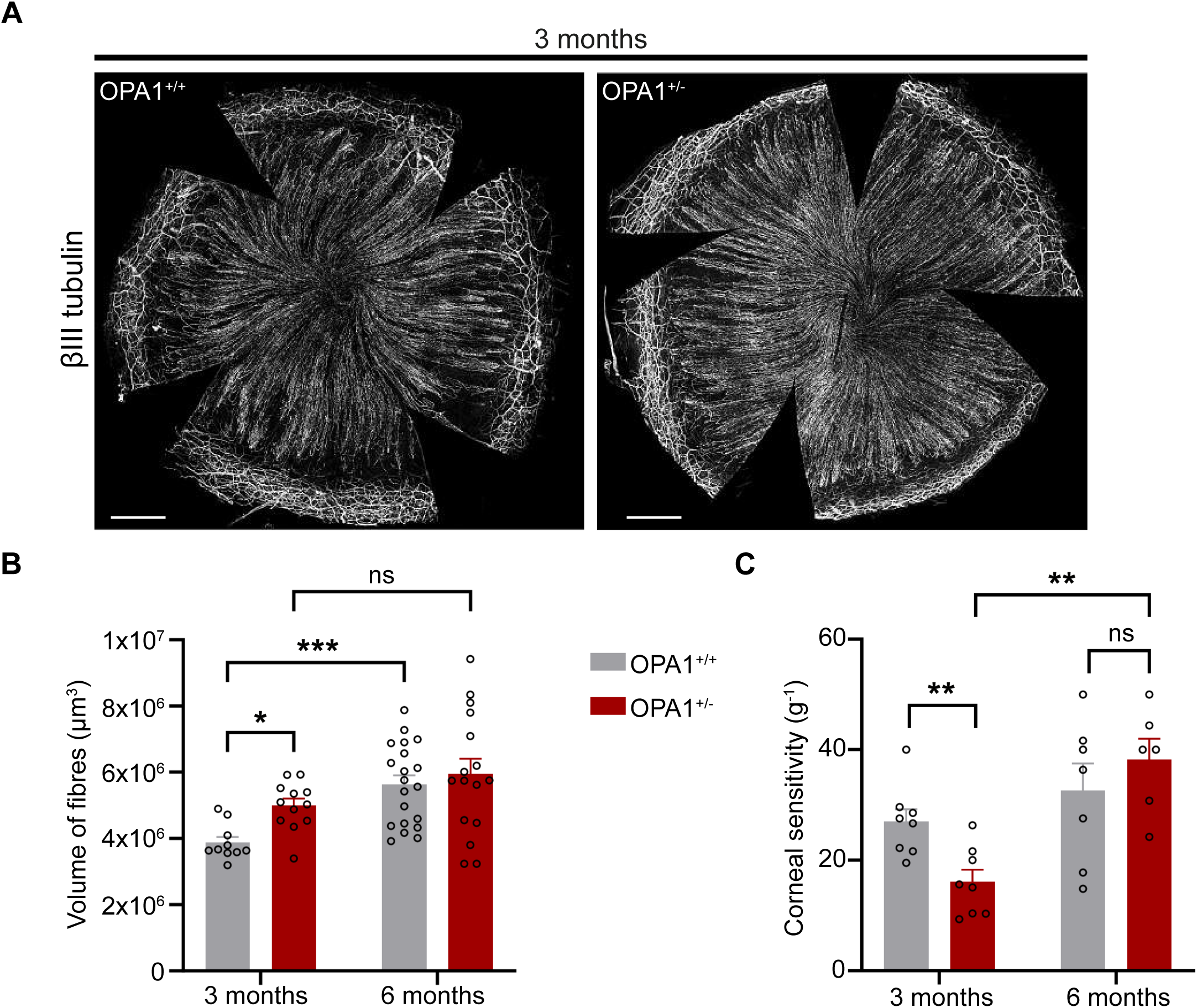
Volume and sensitivity of corneal innervation is impacted in OPA1^+/-^ mice. **(A)** Immunostaining of whole corneas from 3 month-old OPA1^+/-^ and OPA1^+/+^ animal using a pan neuronal antibody (βIII-tubulin) shows a similar pattern of corneal innervation, with fibres extending from periphery to center. Scale bar = 500µm. **(B)** The measurement of corneal innervation highlights a higher volume at 3 months upon the OPA1 mutation, but not at 6 months (n=10-20). Data are represented as mean ±SEM. Statistical anlaysis using two-way ANOVA with uncorrected Fisher’s LSD, and *p<0.05; ***p<0.001. **(C)** The corneal sensitivity measurements assess the low sensitivity upon OPA1 mutation, and the significant gain of sensitivity between 3 and 6 months (n=6-8). Data are represented as mean ±SEM. Statistical anlaysis using unpaired t-test, and **p<0.001.

### OPA1 mutation leads to the modification of corneal nerve identity

Because of the divergence between a higher innervation volume and lower corneal sensitivity, we investigated a possible switch in fibre identity upon mitochondrial dysfunction, which could explain their lower sensitivity transduction. The cornea is innervated by different types of fibres that can be discriminated from each other by the molecules they expressed or secrete (see for review^37^). For instance, the C peptidergic fibres contain calcitonin gene-related peptide (CGRP) and substance P (SP), the cold sensing fibres express transient receptor potential cation channel subfamily M member 8 (TRPM8), and the type A fibres express the neurofilament 200 (NF200). We evaluated the volume of each type of fibres within 3 month-old corneas.

At 3 months of age, we did not measure an impact of OPA1 mutation on the volumes of NF200+, TRPM8+ and CGRP+ fibres. Interestingly, OPA1^+/-^ mice displayed a significantly lower volume of SP+ fibres in the central cornea (Fig.5).

**Figure 5.**
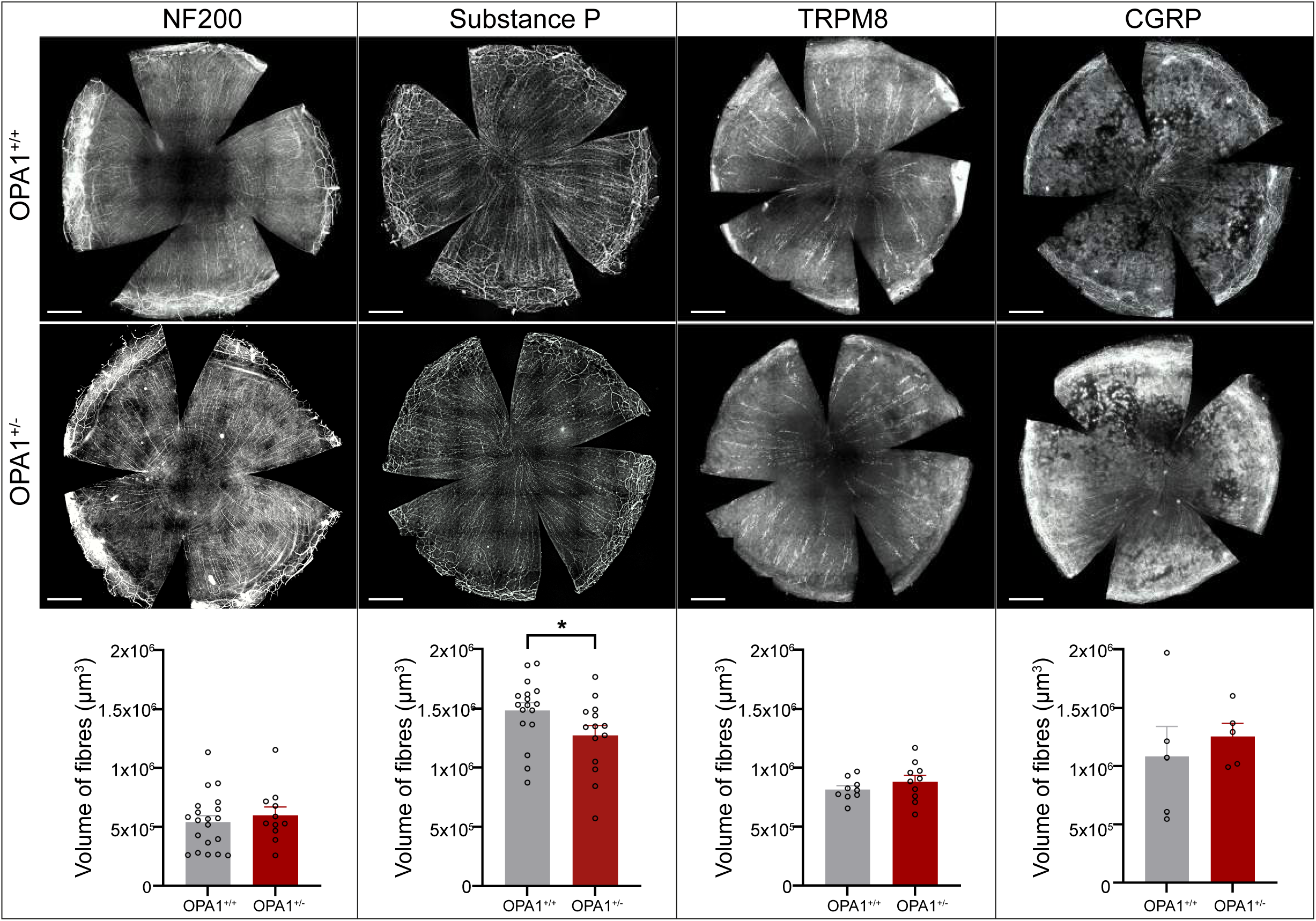
Visualization and volume analysis of the different types of fibres innervating the corneas. The fibres are visualized with anti-NF200, anti-Substance P, anti-TRPM8 and anti-CGRP antibodies. The fibre volumes are then measured for each type of fibres using Imaris software (n=5-20). Volumes of NF200+, TRPM8+ and CGRP+ fibres are not affected upon OPA1 mutation. However, the volume of substance P+ fibres is decreased in OPA1^+/-^ animals. Data are represented as mean ±SEM. Statistical anlaysis using unpaired t-test or Mann Whitney test, and *p<0.05. Scale bar = 500µm.

Because of the modifications of corneal innervation in the context of mitochondrial dysfunction, we investigated the presence of L1 cell adhesion molecule (L1CAM) positive fibres in our model. Among the factors involved in the innervation setup L1CAM is a multifunctional protein, known to be involved in the Sema3A-dependent axone guidance system^38–40^, crucial for corneal innervation^41,42^. We analysed the volume of the L1CAM+ fibres in the central cornea (Fig.6A). Notably, all the nerve fibres of the OPA1^+/+^ cornea seemed to be L1CAM+ at 3 months, as we measured a similar volume for L1CAM+ (Fig.6B) and βIII-tubulin (Fig.4B). Furthermore, we discovered that, unlike βIII-tubulin, the volume of L1CAM+ fibres was lower in OPA1^+/-^ animals at 3 months. Interestingly, at 6 months, the OPA1^+/+^ and OPA1^+/-^ animals had similar volumes of L1CAM+ fibres. Moreover, we did not measure any increase of L1CAM+ fibre volume between 3 and 6 months in neither OPA1^+/+^ and OPA1^+/-^ animals, as we reported with βIII-tubulin (Fig.4B). This observation reflected a change of the surface proteins found on the axons during time.

**Figure 6.**
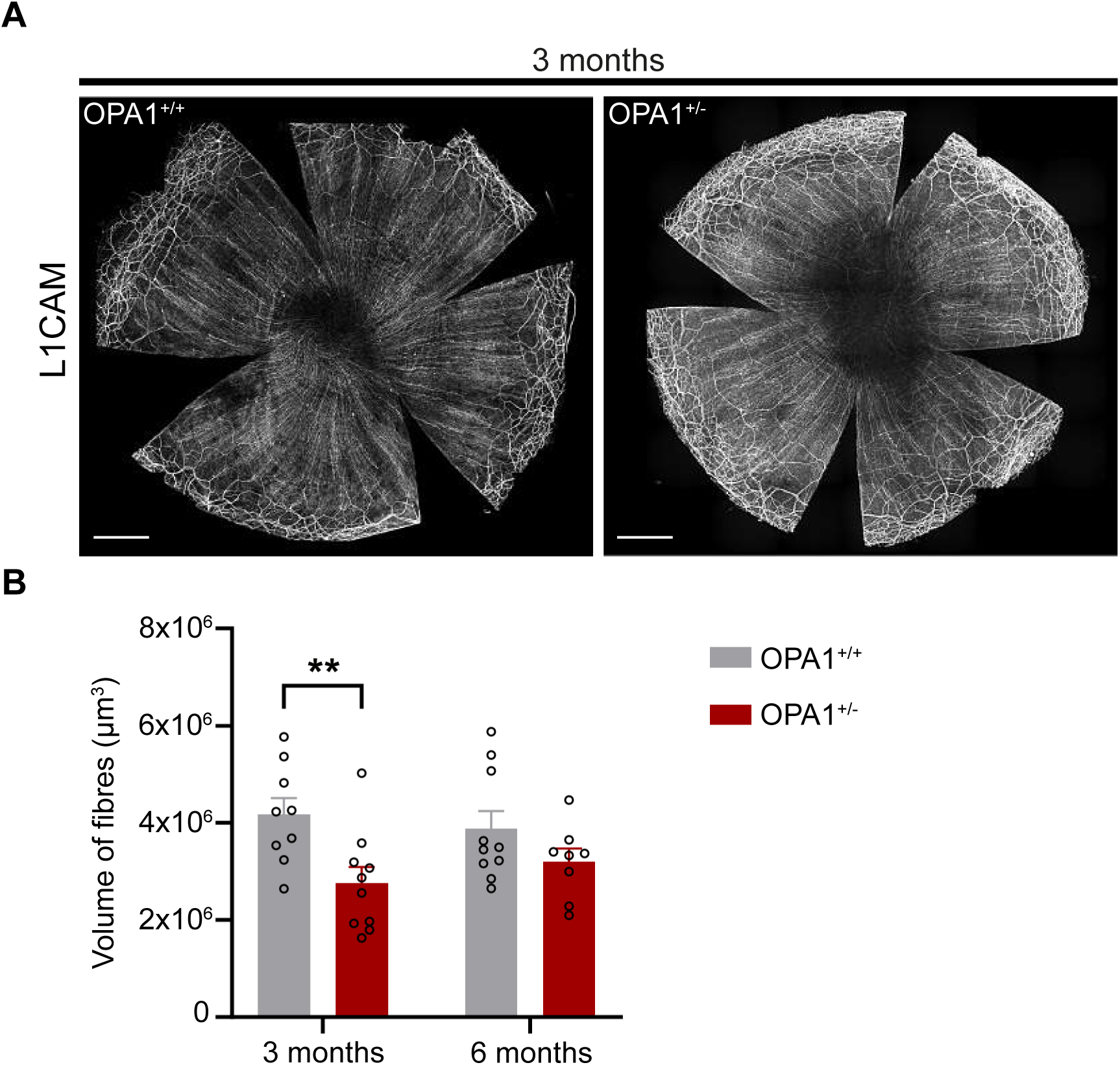
L1CAM+ fibre volume in corneal innervation. **(A)** Whole-mount corneas of OPA1^+/+^ and OPA1^+/-^ animals are stained with anti-L1CAM antibody. The staining shows a normal gross morphology in OPA1^+/+^ and OPA1^+/-^ animals, but a lower staining in the central cornea. **(B)** The measurement of innervation volume reveals that in OPA1^+/-^ mice, L1CAM+ fibres have a lower volume in the central cornea at 3 months, but not at 6 months (n=9-10). Data are represented as mean ±SEM. Statistical anlaysis using 2way ANOVA with uncorrected Fisher’s LSD, and **p<0.01. Scale bar = 500µm.

### OPA1 mutation modifies innervation regeneration dynamics

As the OPA1 mutation impacted corneal innervation in unchallenged situation, we investigated the possible impact of mitochondrial dysfunction on innervation regeneration after corneal abrasion. We injured the cornea using an olphalmic burr to remove the epithelial layer and thereby triggering the healing mechanism^43^. The epithelial wound closure was monitored everyday using fluorescein staining to measure the surface of the epithelial area still unclosed (Fig.7A). Similarly to our previous work have shown^44,45^, the corneal epithelium wound was closed 3 days after abrasion. Moreover, we did not observe a difference in the epithelial closure time upon OPA1 mutation. Then, we assessed the corneal sensitivity during 4 weeks following the epithelial abrasion (Fig.7B). We discovered a significant delay in the loss of corneal sensitivity following the abrasion upon OPA1 mutation. While the lowest sensitivity was measured a week after the abrasion in the OPA1^+/+^ animals, the OPA1^+/-^ mice displayed their lowest corneal sensitivity 2 weeks after the abrasion. Furthermore, we observed a delay in corneal sensitivity recovery upon OPA1 mutation, which was complete 4 weeks after abration in OPA1^+/+^ mice, and was still significantly lower in OPA1^+/-^ animals. Therefore, we investigated the innervation regeneration pattern at 4 weeks post abrasion (WPAb). Two types of reinnervation stood out: either axons started regenerating from the periphery to the centre, similarly to the physiological innervation pattern, we branded this situation as normal, or they emerged from the centre of the cornea, and we branded it as abnormal (Fig.8A). In OPA1^+/+^ mice, more than two third of the corneas had their axons extending from the periphery, whereras for OPA1^+/-^, it occured in less than one-third of the corneas (Fig.8A). Interestingly, we found anarchical growth of axons, forming bushes, regardless the OPA1 status (Fig.8B).

**Figure 7.**
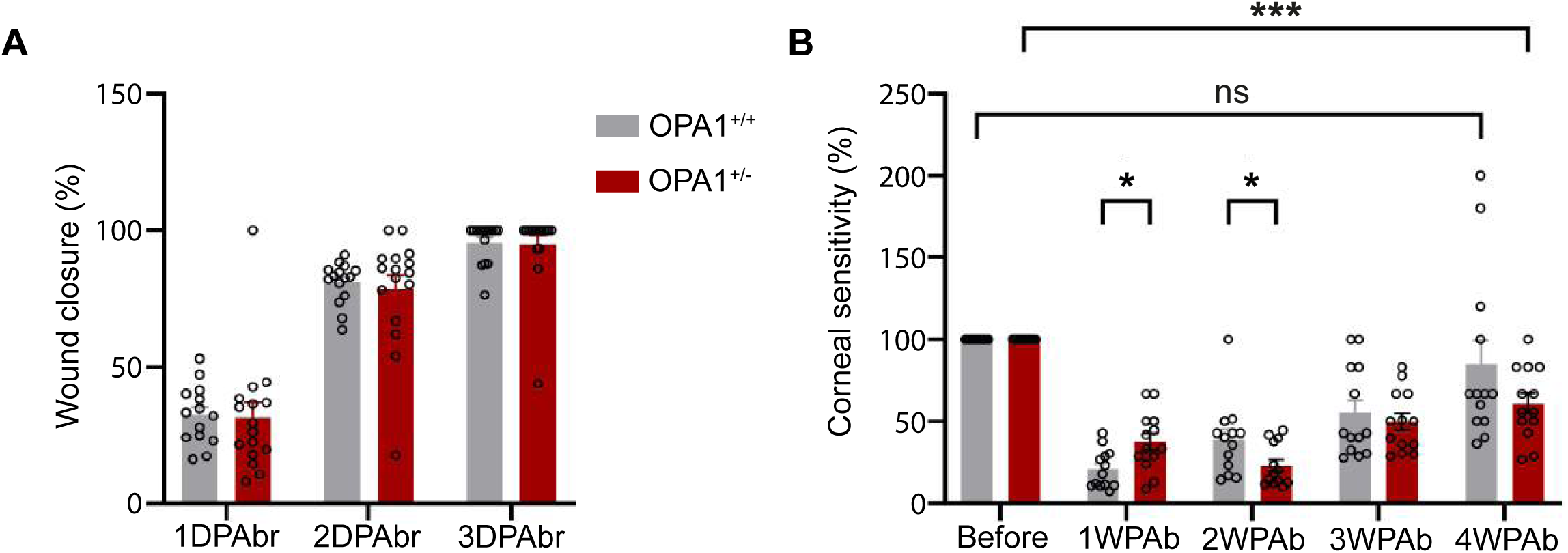
Analysis of the healing properties in OPA1^+/-^ animals. **(A)** Wound closure measurements of the abraded surface reveal no difference in the closure speed between OPA1^+/+^ and OPA1^+/-^ animals (n=14-16). Data are represented as mean ±SEM. Statistical anlaysis using mixed-effects analysis with Sidak’s multiple comparisons test. **(B)** Central corneal sensitivity is monitored up to 4 weeks post-abrasion. The recovery of sensitivity shows a lower decrease of sensitivity at 1WPAbr for the OPA1^+/-^ animals compares to their littermates. At 4WPAbr, OPA1^+/+^ animals recover their sensitivity compared to baseline wherease OPA1^+/-^ animals still have a lower sensitivity (n=13-15). Data are represented as mean ±SEM. Statistical analysis using mixed-effect analysis with Sidak’s multiple comparisons test. *p<0.05 and ***p<0.001. DPAbr: day(s) post abrasion ; WPAbr: week(s) post abrasion. Scale bar = 500µm.

**Figure 8.**
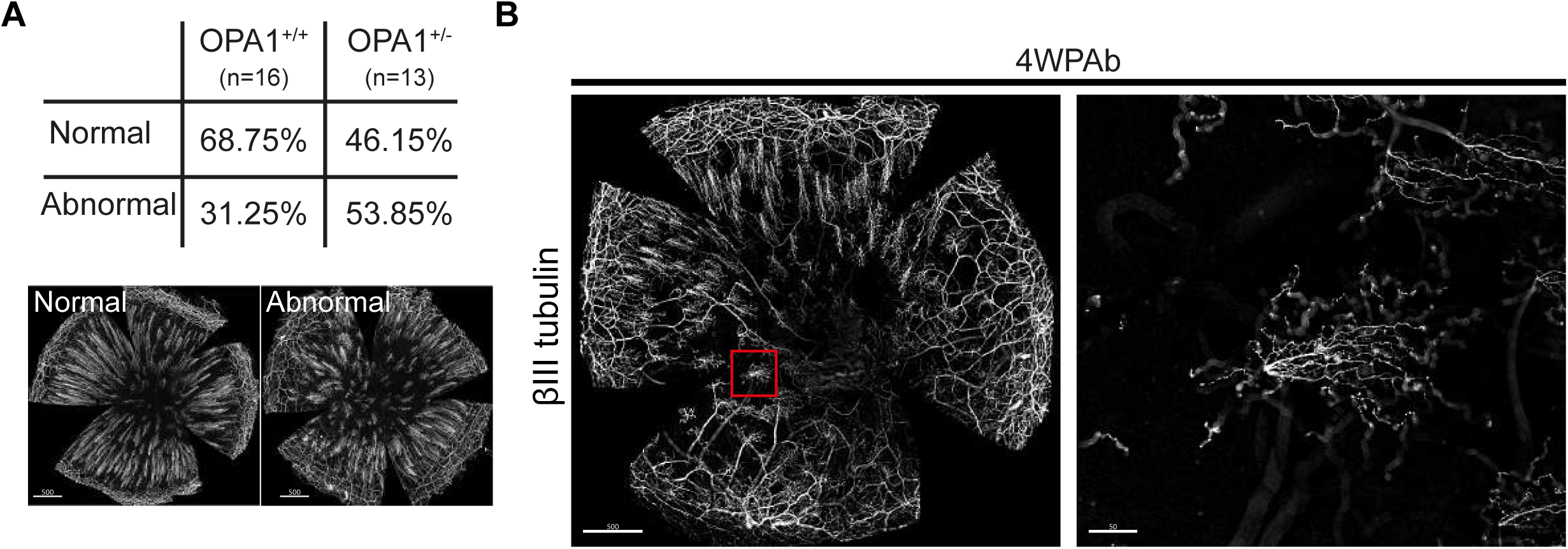
Comparison of the pattern of innervation regeneration. **(A)** The percentage of animals having an abnormal innervation regeneration pattern at 4WPAbr is higher for the OPA1^+/-^ animals (n=13-16). **(B)** Abnormal innervation regeneration in the OPA1^+/-^ central cornea stained with anti βIII- tubulin antibody at 4WPAbr. Red square shows close-up of abnormal innervation regeneration in the OPA1^+/-^ central cornea . WPAbr: week(s) post abrasion. Scale bar = 500µm

As the sensitivity was still significantly lower at 4 weeks after abrasion in OPA1+/- animals, we analyzed the βIII-tubulin+ fibres, SP+ fibres and L1CAM+ fibres recovery at this stage (Fig.9). While the general volume of fibres (βIII-tubulin+) was regenerating slower in OPA1^+/-^ animals, when compared to the OPA1^+/+^, we measured a SP+ fibre regeneration twice as strong in the OPA1^+/-^ animals compared to the OPA1^+/+^ animals. Finally, no difference was detected in the L1CAM+ fibres.

**Figure 9.**
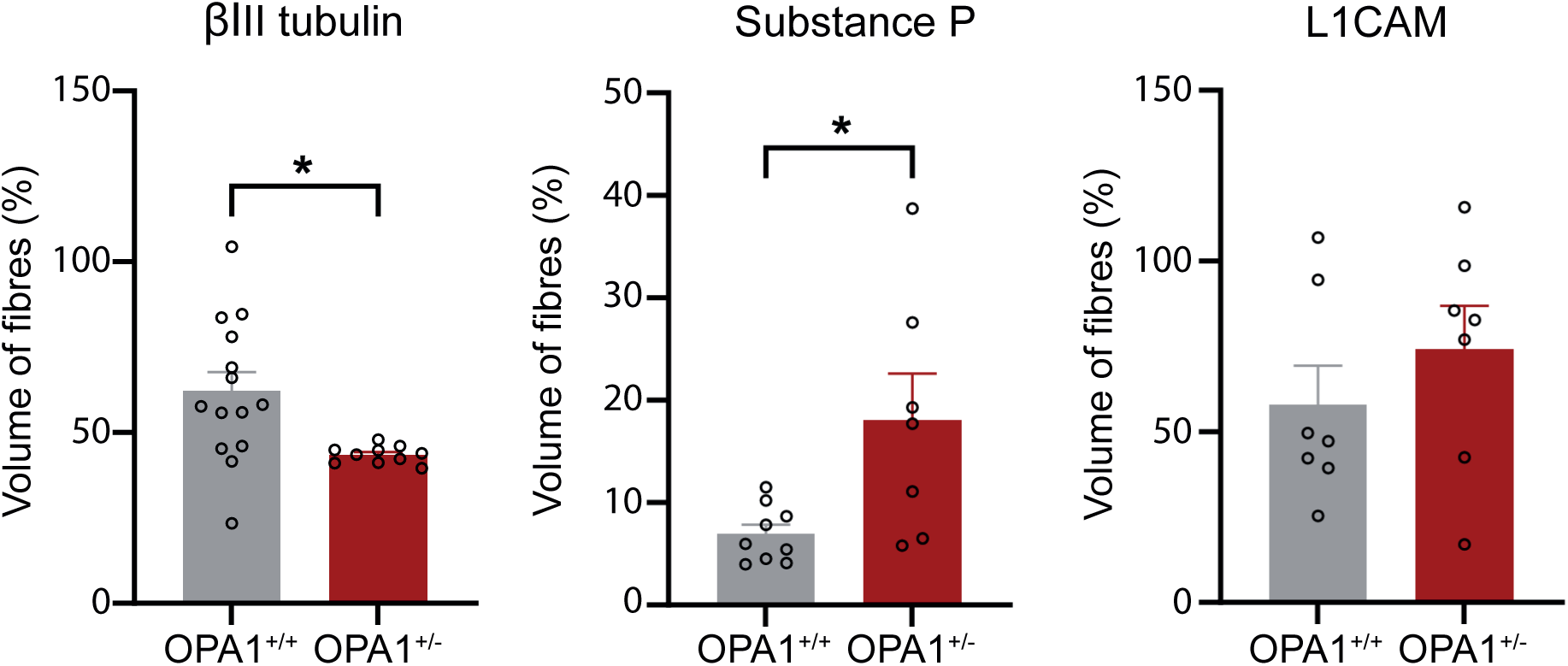
Analysis of the types of fibres reinnervating the cornea at 4WPAb. The volume innervation mesurements indicate that the proportion of fibres reinnervating the cornea at 4WPAb is impacted in OPA1^+/-^ animal regarding βIII tubulin+ fibres and SP+ fibres but not L1CAM+ fibres (n=7-14). Data are represented as mean ±SEM. Statistical anlaysis using unpaired t-test and, *p<0.05. WPAbr: week(s) post abrasion.

## Discussion

Mitochondrial dysfunctions are a large family of pathologies in which metabolism, respiratory chain, and ROS level are impaired. Previous reports have exposed the sensitivity of the cornea to mitochondrial defects, mainly in patients^8,9,12,14,46^. However, the impact of inherited mitochondrial dysfunction on cornea in animal models was still poorly known.

Here, we investigated the impact of the inherited OPA1^delTTAG^ mutation on corneal physiology. The *OPA1* gene encodes a mitochondrial protein essential for the maintenance of mitochondrial integrity and function. Furthermore, patients afflicted with this mutation exhibit optic nerve atrophy, ataxia, and peripheral neuropathy^6^.

The corneal physiology requires a fine-tuned microenvironment. The tear film is part of the ocular surface microenvironement and helps maintaining its homeostasis by providing specific trophic factors to the epithelial surface. Furthermore, the modulation of the tear film composition reflects modification of corneal physiology^45,47,48^. Our proteomic results strongly suggested that the tear film was not affected by the OPA1 mutation. Because a change in tear film composition should reflect a lacrimal gland defect, and/or a subsequent modification of corneal physiology, we hypothesized that in our model any changes with the corneal epithelium would have to be subtle. Therefore, we investigated the transcriptomic signature of the cornea, which has proven before to be a solid indicator of corneal physiology^44,49,50^. As suggested by the unaffected tear film, the transcriptomic signature was largely normal, and all the corneal epithelial identity genes, such as *krt14*, *krt3*, *krt12*, *pax6* were not influenced by OPA1 mutation. Furthermore, our corneal wound healing period was not altered by the OPA1 mutation.

Taken together, these results on corneal resiliency to this mitochondrial defect indicate a possible mitohormesis^51^. This phenomenon is an adaptation to a continuous low-grade mitochondrial stress leading to cells or organs being insensitive to subsequent perturbations. This phenomenon has been recently described in the liver of animals bearing another OPA1 mutation^51^. Therefore, it is possible that an acute OPA1 defect, induced during adulthood, could provoke a stronger corneal defect compared to an inherited mutation for which the whole organ has adapted since its morphogenesis, and found a physiological balance.

Besides the tear film and the epithelium, the sensory innervation is the last component of the corneal microenvironnement. The cornea is composed of a multitude of sensory fibres informing on its well- being. The integration of such information by the central nervous system can lead to the modulation of the tear composition and secretion of proper trophic factors to support the corneal biology. Furthermore, previous studies have shown that the loss of corneal sensitivity was a strong indicator of neurodegenerative disease progression^52–54^. Our results demonstrated that the corneal sensitivity did not decrease with time upon OPA1 mutation. We can conclude that without corneal wounding, there is no development of corneal neuropathy with time.

Our thorough analysis of the corneal innervation revealed that OPA1 mutation induced a more pronounced phenotype in females than in males. This observation is in line with a previous report, based on this specific model, which established that the severity of the optic atrophy is sex- dependent. This has also been confirmed in human where a restrospective analysis reveals a more severe loss of visual acuity and retinal ganglion cell (RGC) degeneration in OPA1 mutated females before the age of 20^55^. Indeed, the mitochondira is an essential part of the steroidogenesis. The production of sex steroid during puberty and lifelong hormonal cycles in females increases the severity of the phenotype comparing to males^56^.

Our results also showed that with age, corneal sensitivity reached a similar level when comparing OPA1^+/+^ and OPA1^+/-^ animals, the mutation only delayed this sensitivity acquisition. As the optimal volume of fibres was acquired faster in OPA1^+/-^ animals, we suggest a decorrelation between the level of innervation and its functionality, most likely supported by an identity plasticity. In pathological contexts, studies have described such phenotypic plasticity, likely promoting corneal wound healing. For example, increased CGRP expression in corneal neurons of a dry eye disease mouse model was demonstrated^57^ and similar increase was seen with TRPM8 axonal expression in an abrasion model^58^. Our staining of differents types of fibres at 3 months showed that SP+ fibres were decreased in OPA1^+/-^ epithelium. SP is a neurotransmitter known to relay intense nociceptive information to the brain^59^. A decrease of SP fibres in the cornea might then explain the decrease of sensitivity seen at 3 months in OPA1^+/-^ mice.

Another key actor of pain sensation is L1CAM molecule. Despite its roles in the mechanism of neuropathic pain^60^, L1CAM has been as well associated with neurite outgrowth^61^, myelination^62^ and regeneration^63^. Our results on the evolution of βIII-tubulin and L1CAM volumes demonstrated that at 3 months in OPA1^+/+^ animals, most of the βIII-tubulin+ axons were L1CAM+, as the ratio of the volumes equal to 1. However, at 6 months this ratio L1CAM volume / βIII-tubulin volume became lower than 1. Interestingly, this ratio was already lower than 1 at 3 months in OPA1 mutant cornea, and was stable from 3 to 6 months. Furthermore, the innervation regeneration after abrasion was slower and disorganized, and the sensitivity was still altered 4 weeks after abrasion in OPA1 mutant mice. We hypothesize that L1CAM is a marker of innervation morphogenesis. As the volume of innervation is already maximum by 3 months in the OPA1^+/-^ animals, the guidance system for the morphogenesis probably happened earlier than in the OPA1^+/+^ animals. By 6 months, the L1CAM volume decreases in our control animals due to the end of the innervation establishment. Furthermore, after abrasion, the L1CAM+ fibres were strongly present to guide the innervation regeneration. Of course, this hypothesis raises 2 critical questions, i) what are the other factors involved in neurite growth? Could the Nrp1 and Sema3A factors be involved in the corneal axon guidance with L1CAM ^64^? ii) what is the corneal innervation maturation process, and fibres identity acquisition? As the corneal sensitivity increased in the OPA1^+/-^ animals from 3 to 6 months without a change in the innervation volume, there is a clear decorrelation between the presence of the innervation and its functionnality. Answering to these questions would unveil knowledge on corneal innervation regeneration in pathophysiological contexts.

The resilience of the corneal biology, and its functionality, upon OPA1 mutation, is outstanding. By developing a mitohormesis, similarly to the liver, the cornea is able to adapt to this mitochondrial dysfunction and find a suitable balance. While the corneal alterations are subtle, the physical harm of the epithelium can trigger a stronger defect by altering the adjusted balance. The factors involved in this balance are still unknown, but discovering them could give new answers on cornea morphogenesis and corneal epithelial identity. They could potentially be therapeutic targets in progressive corneal failure. Finally, while the cornea is overlooked in DOA patients, preserving its balance is important to avoid a sight-threatening corneal failure.

## Material and Methods

### Animals included in this study

The heterozygous OPA1^+/-^ mouse strain^6^ was housed on a 12h dark/light cycle with food and water *ad libitum*. The experiments were conducted on Opa1^+/-^ mice and their Opa1^+/+^ littermate control mice. Animals were euthanized by cervical dislocation.

The animals included in the experiments were approved by the local ethical committee and the Ministère de la Recherche et de l’enseignement Supérieur (authorization 2016080510211993 version2). All the procedures were conducted according to the French regulation for the animal procedure (French decree 2013-118) and with specific European Union guidelines for the protection of animal welfare (Directive 2010/63/EU).

### Cornea abrasions and healing monitoring

Corneal abrasion was performed as previously described. A circular portion of the epithelium of the right eye was removed using and ocular burr (Algerbrush II, reference BR2-5 0.5 mm, The Alger Company, USA) on mice anesthetized with ketamine (80 mg/kg, Imalgene 1000) and xylazine (10 mg/kg, Rompun)(i.p.). The abraded surface was observed using fluorescein solution (1% in PBS, Sigma-Aldrich) under blue cobalt light. After the abrasion, Ocrygel was applied to the eyes and mice were treated with buprenorphine (0.1 mg/kg, Buprecare) (i.p.).

The healing of the abrasion was followed everyday using fluorescein solution on immobilized mice and an image was taken with a camera.

### Sensitivity test

Sensitivity was measured using Von Frey filaments (Bioseb, France, reference bio-VF-M) as previously described ^65^. Filament generate a force between 0.008 and 0.6g when pressed against the centre of the cornea of an immobilized mouse. The weight is increased by changing the filament until an eye-blink reflex is observed. The Von Frey test was performed three consecutive days and values were averaged for each cornea. To reflect the sensitivity, we displayed the value as g^-1^. These experiments were performed by the same experimenter in single-blinded conditions.

### Tissue collection and processing

The mice were euthanized by cervical dislocation. Then the eyes were enucleated by cutting the optic nerve and fixed in 4% paraformaldehyde solution (Antigenfix) for 20 min. After PBS washes, eyes were either immunostained directly or dehydrated 2h in 50% ethanol/PBS and then stored in 70% ethanol/PBS at 4°C.

### Optical coherence tomography (OCT) imaging

Cornea images were acquired *in vivo* with Envisu R2200 spectral domain (SD) OCT apparatus (Leica). The mice were anesthetized with ketamine (80 mg/kg, Imalgene 1000) and xylazine (10 mg/kg, Rompun)(i.p.) and placed on the elevated plateform with a drop of artificial tear on the controlateral eye to avoid desiccation. Images were acquired using the InVivoVue Clinic software (Bioptigen, Inc., Durham, NC) with the 12 mm telecentric lens. The thickness of the layers composing the cornea were measured using the calipers from the software Bioptigen Diver.

### Tear collection

Tear samples were collected successively from both eyes for 1 minute each using 1µL microcapillary glass tubes (Merck, Drummond Scientific, reference P1424-1PAK). The samples from both eyes were pooled in the same tube for each mice, in a total volume of 35µL of Ammonium Bicarbonate (ABC) buffer. The samples were kept at 4°C on ice during the sampling process at the animal facility, and then stored at -80°C for long term storage.

### Total protein concentration and in-solution digestion

Tear samples were thawed at 4°C on ice and the total protein concentration was measured with a NanoDrop One Microvolume UV-Vis Spectrophotometer (Thermo Fisher Scientific, reference ND- ONE-W) by using 2µL of sample. After the protein concentration measurement, the samples were digested directly in-solution. The reduction step was performed by adding a 200mM Dithiothreitol (DTT), 100mM TricHCl (pH 8.5) solution to the samples (DTT final concentration: 20mM). Samples were incubated for 1h at 56°C, 350rpm, protected from light. Then, the alkylation step was done by adding a 300mM 2-Iodoacetamide (IAA), 100mM Tris HCl (pH 8.5) solution to the samples (IAA final concentration: 40mM). Samples were incubated for 30 minutes at 37°C, 350rpm, protected from light. The digestion was achieved by adding 1µg of Trypsin Gold from a 1µg/µL solution, 60µL of 100mM Tris HCl (pH 8.5) and a 200mM DTT, 100mM Tris HCl (pH 8.5) solution (DTT final concentration: 10mM). Samples were incubated overnight at 37°C, 450rpm, protected from light. On the next day, the digestion was stopped by adding formic acid (FA, final concentration: 2%). The samples were then desalted with C18 tips with the AssayMAP Bravo Protein Sample Prep Platform (Agilent Technologies), by using the Peptide Clean-Up program. The loaded volume was set to 108µL and the elution volume was set to 30µL. After the clean-up, the eluted samples were dried at 50°C with an acid-resistant CentriVap centrifugal vacuum concentrator (Labconco) connected to a Centrivap -50°C cold trap (Labconco) and a RZ6 rotary vane pump (Vacuubrand, reference 20698130). The dried samples were temporarily stored at -20°C.

### Mass spectrometry analysis

The samples were resuspended in 10µL phase A of Q-TOF (2% ACN, 0.1% FA in water HPLC grade) in mass spectrometry polypropylene vials (Agilent technologies, reference 5190-3155). The samples (1µL out of 10µL) were analyzed online by using the NanoElute® High-Performance Liquid Chromatography (HPLC) (Bruker) coupled with the Impact II Quadrupole Time-Of-Flight (Q-TOF) mass spectrometer (Bruker). Each sample was analyzed in triplicate. The peptides were separated on a C18 column (FORTY, reference 1842619 from Bruker; 75µm x 400mm; Acclaim Pepmap RSLC, C18, 1.9µm, 120Å) following a gradient of 2-15% phase B in 55 minutes, 15-25% in 25 minutes, 25- 35% in 10 minutes and 35-95% in 10 minutes (phase A = 2% ACN, 0.1% FA in water HPLC grade; phase B = 0.1 % FA in 90% ACN). The column was kept at 50°C and the flow rate was set to 300 nL/min. Water, acetonitrile (ACN), formic acid (FA) were all HPLC grade and purchased from Biosolve (Dieuze, France).

### Data processing and statistical analysis after Mass-Spectrometry analysis

Protein identification was performed using MaxQuant software (1.6.17.0 version). The parameters were as follows: the enzyme used for protein cleavage was the trypsin, the mass tolerance was set to 10ppm for the precursor ions and to 0.005 Da for the MS/MS. The murine proteome from the Uniprot database was used (April 2022 version). The carbamidomethylation of the cysteine was programmed as a fixed modification, the oxidation of the methionine and the deamidation of the asparagine and glutamine were programmed as variable modifications. The maximum number of modifications for one peptide was set to 5.

Data processing was performed using Perseus software (1.6.15.0 version). Proteins considered as contaminants and redundant proteins were deleted. Data were grouped following the age (3.5 months or 6 months) and the genotype (Opa1^+/-^ or Opa1^+/+^). The values obtained for each protein were transformed by using the log2(x) formula. For each time point, the proteins must be found in the 3 biological replicates of each genotype in order to be conserved. Data were then normalized in order to follow a normal distribution.

Correlations between the protein composition and the parameters (age or genotype) were analyzed following a distance calculated with the Pearson correlation coefficient. Differences in the protein composition among groups were tested using the Student t-test, with a permutated-based false discovery rate (FDR) of 0.05 (p-value < 0.05 was considered statistically significant).

### RNA-sequencing

Total RNA was extracted using Trizol reagent and purified afterwards with Qiagen RNeasy kit following the manufacturer’s procedure. Total RNA concentrations were measured with Qubit and quality was assessed using RNA ScreenTape Assay (Agilent TapeStation 4200). 500 ng of good quality (RIN >8.8) total RNA was ribo-depleted using Illumina’s mammalian Ribo-Zero Magnetic Gold Kit HMN/Mouse/Rat (MRZG12324). Library preparation was completed on ribo-depleted RNA using the NEBNext Ultra Directional RNA Library prep kit (NEB, #E7420L) using 8 cycles of PCR amplification and indexed using single (i7) indexing. Indexed library preps from each sample were then pooled and sequenced with 75SE reads at a pool concentration of 1.4 pM on the NextSeq 500 using a NextSeq High Output 75 cycle flow cell (Illumina).

### RT-qPCR

Total RNA was extracted from samples containing each 4 mouse corneas. Tissue grinding was performed with 1 ml TRIzol reagent (Life Technologies) using CKMix lysing kit and Precellys homogenizer (Bertin Technologies) followed by RNA purification and DNase treatment with the Nucleospin RNA Kit (Macherey Nagel). RNA samples were quantified using the NanodropOne spectrophotometer (ThermoFisher Scientific) and the integrity of the RNAs was verified using the Agilent 2100 bioanalyzer with the RNA 6000 Nano kit (Agilent Technologies). 1 µg of total RNA was reverse transcribed in a 20 µL μnal reaction volume using the High-Capacity Reverse Transcription kit (Applied Biosystems) using random primers and the presence of RNase inhibitor following the manufacturer’s instructions. Quantitative PCR experiments were performed with the QuantStudio 12K Flex Real-Time PCR System (Life Technologies) using a TaqMan detection protocol. TaqMan assays (ThermoFisher Scientific) were selected for each target and reference gene (Table 2). For qPCR reactions, 3 ng of cDNA were mixed with TaqMan FastAdvance Master Mix and TaqMan assay in a final volume of 10 µl, loaded on 384 well microplates and submitted to 40 cycles of PCR (50°C/2 min; (95°C/1 sec; 60°C/20 sec) X40). Negative controls without the reverse transcriptase were introduced to verify the absence of genomic DNA contaminants. Each sample measurement was made in duplicate. Determined Ct values were then exploited for further analysis. To normalize the data 3 reference genes were also added in the experiments. The most stable reference genes were selected by GenEx software (MultiD) and the geometric mean of the 2 most stable genes (Hprt1 and Ppia) was used to normalize the data. The determination of the relative gene expression ratio was achieved using the ΔΔCt method.

### Immunostaing of whole corneas

The eyes, where corneal innervation density was measured, were rehydrated for 2h in 50% ethanol/PBS at room temperature. After 2 PBS incubations of 10min, corneas were dissected and then blocked and permeabilized in FSG 2,5% (Sigma-Aldrich, reference G7765) and goat serum (GS) 5% (Thermo Fisher Scientific, reference 16210064) in 0,5 % PBS/Triton X-100 at room temperature. The corneas were then incubated with primary and secondary antibodies (Table 1) overnight at 4°C in FSG 2,5% and GS 5% in 0,1 % PBS/Triton X-100 and rinsed 3 times for 1 hour in 0,1% PBS/Triton X-100. Nuclei were stained with BioTracker NIR694 (Merck, reference SCT118) or Hoechst Hoechst 33342 (Thermofisher Scientific, reference H3570) for 10 min and rinsed in PBS. The eyes collected to stain the different types of nerve fibres in the cornea were not dehydrated. Corneas were directly dissected, blocked and permeabilized in FSG 2,5% and GS 5% in 2% PBS/Triton X-100. Cornea were then incubated with primary and secondary antibodies (Table 1) overnight in FSG 2,5% and GS 5% in 1% PBS/Triton X-100. Cornea were washes 3 times for 1 hour in 1% PBS/Triton X-100. Nuclei were stain using Hoechst 33342 (Thermofisher Scientific, reference H3570) for 10 min.

**Table 1.**

| **Antibody name** | **Host** | **Vendor/Reference** | **Dilution** |
| --- | --- | --- | --- |
| Anti-beta III Tubulin | Rabbit | Abcam / ab18207 | 1/1000 |
| Anti-NFH (Neurofilament Heavy Chain) Antibody | Chicken | Aves labs / AB_2313552 | 1/500 |
| Calcitonin Gene-Related Peptide 1 (alpha-CGRP) | Rabbit | BMA Biomedicals / T-4032 | 1/500 |
| Anticorps Substance P (NC1/34HL) | Rat | Santa Cruz / sc-21715 | 1/500 |
| Anti-TRPM8 (extracellular) Antibody | Rabbit | Alomone labs / ACC-049 | 1/500 |
| Anti-Neural Cell Adhesion Molecule L1 Antibody, clone 324 | Rat | Merck / MAB5272 | 1/500 |
| Goat Anti-Rat IgG H&L (Alexa Fluor® 647) preadsorbed | Goat | Abcam / ab150167 | 1/500 |
| Anti-Rabbit IgG (H+L) Cross-Adsorbed Secondary Antibody, Alexa Fluor 488 | Goat | ThermoFisher / A-11008 | 1/500 |
| Anti-Chicken IgY H&L (Alexa Fluor® 568) preadsorbed | Goat | Abcam / ab175711 | 1/500 |

**Table 2.**

| **Gene** | **ID** |
| --- | --- |
| Adamts1 | Mm01344169_m1 |
| Adgrf2 | Mm01281279_m1 |
| Alox15 | Mm00507789_m1 |
| Arntl | Mm00500223_m1 |
| Ciart | Mm01255905_g1 |
| Clmp | Mm00505221_m1 |
| Cpeb1 | Mm01314928_m1 |
| Nxph4 | Mm00806440_m1 |

Four incisions were performed on all the corneas that were then mounted in Vectashield medium (Vector laboratories, H-1000), with the endothelium facing the slide.

### Imaging and processing

The image acquisition was performed as previously described.^65^ Whole-cornea images were acquired on a Leica Thunder Imager Tissue microscope using the navigator module with the large volume computational clearing (LVCC) process. Images were obtained with LAS X software (3.7.4) using a 20X/0.55 objective and processed with Imaris Bitplane software (version 9.8.0). The same parameters were used to acquired and processed all of the images from a single panel. To analyse the healing of the abrasion, the area of the abraded surfaces was measured using Fiji *measurement* plug-in (FIJI (RRID:SCR_002285)).

### Innervation volume measurement

50% centred circle was draw with FIJI (RRID:SCR_002285) using the *measurement* plug-in to measure the diameter of each cornea twice and use the mean to select the radius of the circle for each cornea. A 50% circular crop of each cornea was obtain by using the *crop* plug-in followed by the *clear outside* plug-in.

The 50% crop images were then converted into *.ims* format on Imaris converter. Innervation density has been measured using the tool surface on Imaris Bitplane software (version 9.8.0). Once the surface has been established, the data regarding the volume in µm^3^ were retrieved to do the statistical analysis.

### Statistical analysis

GraphPad Prism software was used to analyse the data (version 10.1.2). Data are expressed as mean ±SEM and adjusted p < 0.05 was considered significant. Statistical analysis comparing mean values were tested using two-way ANOVA with Sidak’s multiple comparisons test, Benjamini- Hochberg p-value correction, . Statistical anlaysis using two-way ANOVA with uncorrected Fisher’s least significant difference (LSD), unpaired t-test and Mann Whitney test

## Author contributions

Conceptualization, L.M and F.M; methodology, L.M, N.F; formal analysis, L.M and F.M.; investigation, L.M., N.F., S.P., M.G., S.S., A.C., M.Q., L.F. and J.V.; data analysis L.M. and F.M.; writing-original draft, L.M. and F.M.; writing – review and editing, all of the authors; project administration and funding acquisition, F.M.

## Acknowledgments

This research was supported by ATIP-Avenir program, Inserm, the Région Occitanie, ANR (CE17 - Project TeFiCoPa), FRM (REP202110014140), the Fondation de France (00143435 / WB-2023- 48237), CBS2 Doctoral School, University of Montpellier. The RNA-Sequencing service was provided by the Biomedicum Functional Genomics Unit at the Helsinki Institute of Life Science and Biocenter Finland at the University of Helsinki. The RT-qPCR service was provided by the Dr Eric Jacquet at the QPCR plateform from Réseau des plateformes de Génomique Paris-Saclay at the Institut de Chimie des Substances Naturelles at Gif-sur-Yvette. We thank the MRI-DBS imaging facility, member of the France-BioImaging national infrastructure supported by the French National Research Agency (ANR-10-INBS-04, «Investments for the future»).

## Supplemental Material

**Supplementary Figure 1.**
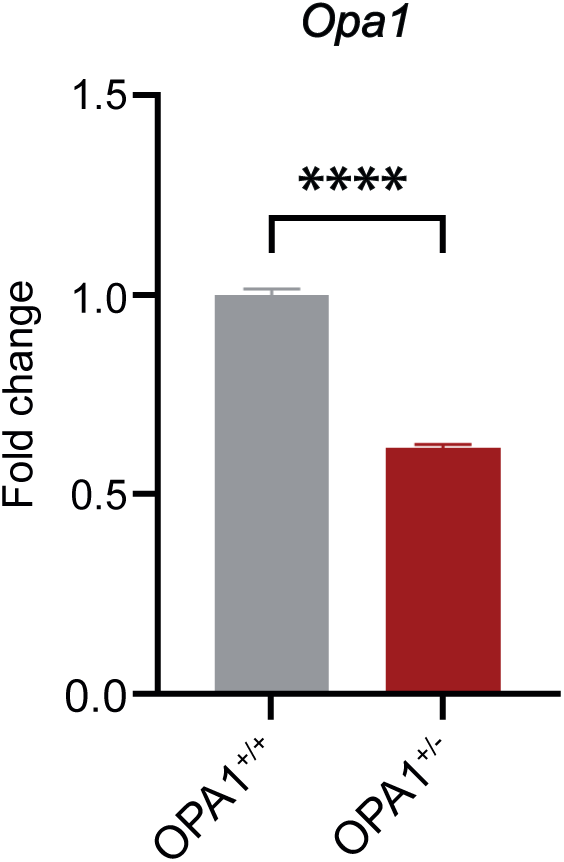
RTqPCR analysis of OPA1 in the cornea. The OPA1 is validated. Data are represented as mean ±SEM. Statistical anlaysis using unpaired t-test, and **p<0.01.

**Supplementary Figure 2.**
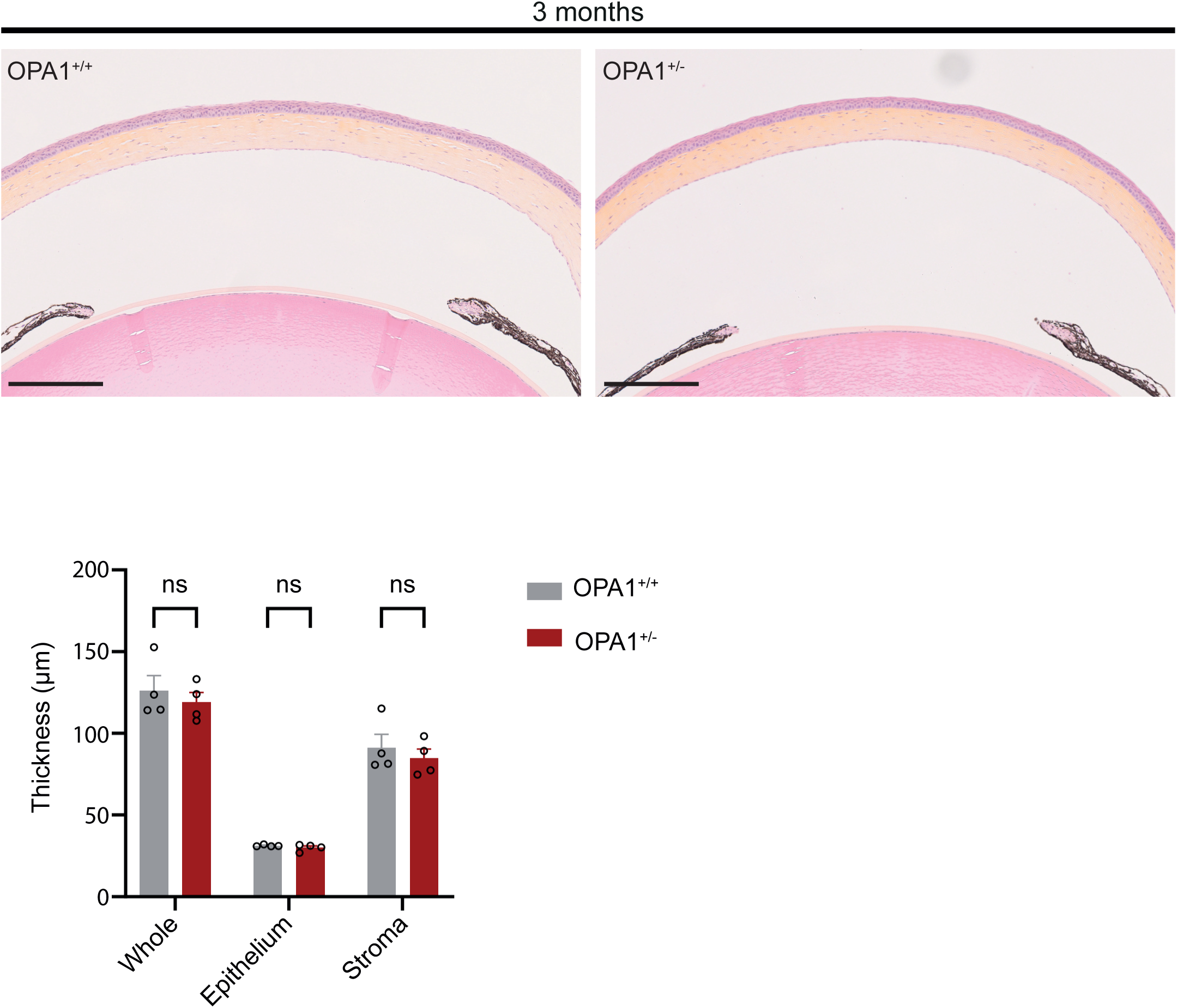
Volume of corneal innervation in OPA1^+/+^ and OPA1^+/-^ male animals. Analysis of the volume of innervation in the central cornea of male animal indicates no difference at 3 months. Statistical analysis using unpaired t-test and ns, p>0.05.

**Supplementary Figure 3.**
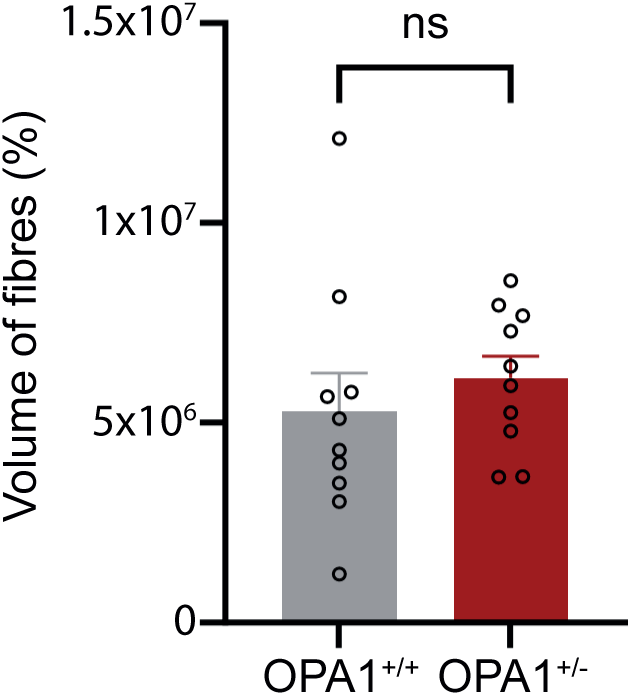
Cornea histological analysis shows a maintained morphology in OPA1^+/-^ animals. Hematoxylin-eosin-safran staining was performed to assess the overall morphology of the cornea. The OPA1 mutation does not impaired the morphology of the corneal layers. Scale bar = 250µm.

